# The Chromosome Periphery is an Essential Compartment of Oocyte Chromosomes

**DOI:** 10.1101/2024.12.20.629576

**Authors:** Eva L Simpson, Ben Wetherall, Aleksandra Byrska, Liam P Cheeseman, Tania Mendonca, Xiaomeng Xing, Alison J Beckett, Helder Maiato, Alexandra Sarginson, Ian A Prior, Geraldine M Hartshorne, Andrew D McAinsh, Suzanne Madgwick, Daniel G. Booth

**Affiliations:** Biodiscovery Institute, Queen’s Medical Centre, University of Nottingham, Nottingham, NG7 2UH; Newcastle University Biosciences Institute, Faculty of Medical Sciences, Newcastle University, Newcastle upon Tyne, NE2 4HH, UK; Biomedical EM Unit, Liverpool Shared Research Facilities, University of Liverpool, Liverpool, UK, L69 3BX; i3S, Instituto de Investigação e Inovação em Saúde, University of Porto, Rua Alfredo Allen, 208, 4200–135 Porto, Portugal; Molecular & Clinical Cancer Medicine, Institute of Systems, Molecular and Integrative Biology, University of Liverpool, Liverpool, UK, L69 3BX; Warwick Bio-Medical Sciences and Centre for Mechanochemical Cell Biology, Warwick Medical School, University of Warwick, Coventry CV4 7AL, UK; University Hospitals Coventry and Warwickshire NHS Trust, Coventry CV2 2DX, UK

## Abstract

Chromosomes and their constituent compartments are core elements of the cell division machinery. In mammalian oocytes, defects in the structure or composition of chromosomes are a leading cause of aberrant or failed meiosis, and by extension, can cause miscarriage and infertility. The underlying mechanisms are poorly understood, but are critical for development of novel diagnostics and treatments.

The chromosome periphery, the least understood chromosome compartment, has recently emerged as an essential component of mitotic chromosomes and somatic cell division. However, in female meiosis it remains completely unexplored.

This study provides the first comprehensive survey of a meiotic chromosome periphery compartment in both human and mouse oocytes. Using a combination of time-lapse microscopy, super-resolution imaging and 3DCLEM we show that removing the chromosome periphery, via Ki67 depletion, has substantial negative impact on chromosome structure, spatial organization and positional awareness, with many oocytes stalling and arresting in meiosis I. Importantly, we also reveal key differences between the mitotic and oocyte meiotic chromosome periphery compartments, most remarkably in the retention of Ki67 in the oocyte through anaphase I, where unwanted chromosomes are stripped of Ki67 before being ejected from the oocyte.

This work presents the discovery of an exciting new pathway operating during female meiosis and a provides a platform for future work exploring the meiotic chromosome periphery for therapeutic vulnerabilities.

## INTRODUCTION

Accurate cell division is essential for all life and relies on the timely organisation of multiple chromosome compartments. Broadly, there are six major regions/compartments of chromosomes. Five of these; nucleosomes/chromatin, telomeres, kinetochores, centromeres, and the scaffold, have been studied extensively, and all linked to highly aberrant cell division, if defective. In the oocytes of mammals, defects in chromosome structure/composition are a major cause of aneuploidy^1^. Indeed, for largely unknown reasons, up to half of human oocytes divide with segregation errors. The outcomes of this defect can be devastating, frequently leading to, infertility, miscarriage and birth defects^2,3^ But aneuploidy isn’t the only trigger of oocyte failure, there are a substantial number of other, even lesser understood, sources of failure. Many of these oocytes stall, preceding a division attempt – with a key phenotype being radical chromosome rearrangements, such as chromosome “clumping”^4^. These oocytes are incompatible with life through natural pregnancy or/and are frequent source of loss through assisted reproductive therapies (ARTs) in clinic. The molecular mechanisms underpinning these events are diverse and unclear, but are important because they correlate with increasing maternal age^3,5–8^ and with an ageing population and socio-economic changes that advance maternal age, these are 21^st^ century challenges that need to be confronted. Thus, there is clearly an unmet need to investigate new pathways of both *healthy* and *unhealthy* female meiotic division.

We address this by performing the first characterization of the least understood chromosome compartment, the chromosome periphery (CP), specifically in oocytes.

To date, almost all work on the CP, an enigmatic sheath that covers the outer surface of condensed chromosomes, has focused during somatic cell division. Indeed, this field has recently been reinvigorated following the discovery that Ki67, the prominent cancer proliferation marker, is the chief organizer of the CP^9^, providing the first opportunity to probe CP biology since its discovery in 1882^10^. As a result, there is now compelling evidence that the CP performs multiple critical roles, including; supporting chromosome individualization (preventing chromatin “clumping”)^11,12^ and clustering during late mitosis^13^, the symmetrical distribution of nucleolar material between daughter cells^9,14^, the maintenance of chromosome architecture, either directly^15^ or via organization of its epigenetic landscape^16^ and finally, as a protector against DNA damage^17^.

The precise details of its function are still to be fully defined, but there is now wide consensus that the mitotic CP should be accepted, like the five other regions, as an essential compartment of mitotic chromosomes^18–20^. Importantly, even though the Ki67 antibody was generated in 1983^21^, in the interim, a definitive Ki67 or CP distribution throughout the key stages of oocyte maturation remained elusive^22^.

Should one exist, it is far from certain that it would be a spatial and functional replica of its mitotic counterpart. For example, why would a function in the equal distribution of nucleolar material benefit a cell that does not reform a nucleolus between divisions? Or, whilst mitotic chromosome individualization might be important for the confined space of somatic cell division, why would this concern oocytes, that have ample volume for free movement?

This raises four important questions. In the event it exists, could an oocyte chromosome periphery be essential to meiotic chromosomes? If so, does it contribute to successful meiotic progression, and is this akin to mitotic functions? Finally, could an understanding of the meiotic CP be of tractable clinical benefit?

Here, we have determined that Ki67 has a novel function during female meiosis and is essential for meiotic progression. We consider that this may be linked to a role in the maintenance of chromosome morphology or chromosome individualisation and speculate of underpinning mechanisms relating to meiotic drive and asymmetric cell division. Importantly, the abundance of Ki67 decorating chromosomes varies significantly between oocytes. Exaggerating this by depletion experiments demonstrates how in oocytes with lower Ki67, chromosomes lose spatial awareness over the volume of the oocyte, ultimately leading to chromosome clumping and a failure to mature – which is a frequent and poorly characterised event, in human subfertility and IVF failure.

## METHODS

### Mouse Oocyte Methods

#### Mouse oocyte collection and culture

Six-to-10-week-old female CD1 mice (Charles River) were used for oocyte collection. All aspects of animal handling were performed in accordance with ethics approved by the UK Home Office Animals Scientific Procedures Act 1986. The project did not require Home Office Licensing, and accordingly animal use was approved and governed by Newcastle University’s Comparative Biology Centre Ethics Committee – AWERB approval reference 663. Prophase-arrested GV oocytes were collected from ovaries punctured by sterile needle and stripped of cumulus cells mechanically by glass pipette. For handling, microinjection, and imaging oocytes were cultured in 37°C M2 medium (Sigma-Aldrich) under mineral oil (Sigma-Aldrich). Where necessary, media was supplemented with 30 nM 3-isobutyl-1-methylxanthine (IBMX) (Sigma-Aldrich) to maintain cell-cycle arrest at prophase I. Oocytes were only collected that appeared visually competent with minimal granularity, a centred germinal vesicle, and the presence of perivitelline space. For each experiment both the control and treatment groups were sourced randomly from the same pool of oocytes, originating from a minimum of two animals.

### Knockdown of Ki67 using morpholinos

A morpholino oligo was used to target and knockdown endogenous Ki67 mRNA in mouse oocytes (sequence AGAAGCCTCTCGGTGAAGCTCGCA, designed and produced by Gene Tools). The Morpholino was used per the manufacturer’s instructions, at a concentration of 1mM in water, and heating for 5 minutes at 65°C prior to use.

Microinjection was performed on the heated stage of an Olympus IX71 inverted epifluorescent microscope, under the culture conditions stated above. Microinjection needles were fabricated from filamented glass capillaries (Harvard Apparatus) using a Model P-97 Flaming/Brown micropipette puller (Sutter Instruments). Morpholino delivery was achieved using a PV830 Pneumatic PicoPump and MICRO-ePore (World Precision Instruments). Injection volume was estimated by ooplasm displacement and was typically in the range of 0.25% cell volume.

### Fixed cell imaging microscopy

For Ki67 localisation experiments, mouse oocytes were attached to the bottom of 8-well chamber slides (Falcon) using 0.2 μL droplets of Cell-Tak (Corning). Oocytes were briefly rinsed with warm PBS-Tween (0.1%) (Sigma-Aldrich) before being fixed in PBS containing 1.6% formaldehyde (Thermo Scientific) and 0.1% Triton X-100 (Sigma-Aldrich) at room temperature for 30 minutes. Fixed oocytes were then further permeabilised by incubation at 4°C overnight in 0.1% Triton X-100 PBS, and stored in PBS-Tween (0.1%) for 48-72 hours. Blocking was performed at room temperature for 2 hours with 3% BSA (Sigma-Aldrich) PBS-Tween. Ki67 Monoclonal Antibody (SP6) (Invitrogen MA5-14520) was used at 1:250 and incubated overnight at 4°C. Goat anti-rabbit Alexa Fluor 488 (Invitrogen A-11008) secondary antibody was used at 1:1000. SiR-DNA was added 1:1000 during secondary antibody incubation to counterstain chromosomes. In order to preserve 3D structure, the oocytes were not mounted in a mounting medium, but rather sealed under a coverslip in PBS and imaged immediately. Ki67 localisation imaging was performed using a Zeiss LSM 800 confocal microscope with Zen Blue software (Zeiss). Imaging was conducted with a 63x oil immersion objective (NA 1.40) with a slice-thickness of 0.23 – 0.27 µm.

### Live imaging microscopy

Live fluorescent imaging tracking meiosis I progression with SiR-DNA was performed at 37°C on a heated stage using either a 20x air lens on a DMi8 widefield microscope with LAS X software (Leica Microsystems), or a 63x water immersion lens on an LSM 880 confocal microscope in AiryscanFAST mode with Zen Black software (Zeiss). Widefield images were captured in approximately 10 focal planes every 10 minutes, while high-resolution confocal imaging to produce 132-slice Z-stacks (slice thickness 0.42 µm) was performed with 30-minute intervals. Control and Ki67 depleted oocytes were stained with SiR-DNA, incubated to resume meiosis and followed by live-imaging for ∼16 hours. Scoring criteria was as follows: Stalled; oocytes did not progress to anaphase within the time window of live imaging (i.e. 16 hours). Typically this meant chromosomes “clumped” into a single contiguous mass, with very little movement. Misaligned: chromosomes failed to congress and align in a typical metaphase arrangement. Mis-segregated: Only gross, readily observable errors were scored, including: ejection of full chromosome set into PB, formation of PB but retention of full chromosome set, clear lagging chromosomes in anaphase or chromosome bridges.

### Knockdown of Ki67 using Trim-Away

#### In vitro mRNA transcription

Messenger RNA transcription was performed using pGEMHE-Trim21 (kind gift from Dean Clift). The plasmid was linearised, and capped mRNA was synthesized using the T7 ARCA mRNA kit (New England Biolabs) according to the user’s manual and resuspended in water. The final concentration of mRNA in the injection needle was 400 ng/µl.

#### Trim-Away experiments

All mice used for Trim-Away experiments were maintained in a specific pathogen-free environment according to the Portuguese animal welfare authority (Direcção Geral de Alimentação e Veterinária) regulations and the guidelines of the Instituto de Investigação e Inovação em Saúde animal facility. Oocytes were freshly collected from female FVB/N mice 8 to 12 weeks old. Oocytes were maintained in M2 medium supplemented with 250 µM dbcAMP under paraffin oil in a CO2-free incubator at 37°C. After extraction from the ovary, oocytes were microinjected with mRNA encoding for Trim21, and incubated for 3 hours to allow for mRNA expression. Oocytes were then injected with anti-Ki67 antibody or control IgG, and immediately released into unsupplemented M2 medium overnight (12-16 hours). The oocytes were then fixed in 4% paraformaldehyde in PBS for 30 minutes, permeabilized with 0.5% Triton X-100 in PBS for 30 minutes, and blocked in PBS containing 3% BSA and 0.1% Triton X-100 for 1 hour. Oocytes were then sequentially incubated with primary and secondary antibodies for 2 hours each at room temperature. Oocytes were then transferred to PBS supplemented with SiR-DNA (500 nM, Spirochrome) under paraffin oil, and imaged on an Abberior ‘Expert Line’ gated-STED microscope on confocal mode (without STED beam), equipped with a Nikon Lambda Plan-Apo 1.4 NA 60x objective lens.

### 3DCLEM

To identify meiotic stages and localize chromosomal positions within oocytes, cells were seeded onto glass-bottom dishes (MatTek Corporation) pre-coated with CellTak (Corning). CellTak was allowed to dry completely before use. Oocytes were then placed on the coated dishes and allowed to settle to permit to adherence. Following adhesion, cells were fixed in a solution of 2% paraformaldehyde (Thermo Fisher Scientific) and 2% glutaraldehyde (Sigma-Aldrich) in 0.1 M sodium cacodylate buffer, supplemented with DAPI (1:1000) for nuclear staining. Fixation was carried out for 1 hour, after which cells were covered with phosphate-buffered saline (PBS) and stored at 4°C until imaging.

High-resolution images were acquired using a Zeiss LSM 900 microscope equipped with AiryScan 2. Imaging was conducted with a 100x oil immersion objective (numerical aperture 1.518) using a UV filter (405 nm excitation). The Zen Blue software platform was utilized for image acquisition. Z-stacks were captured with a slice thickness of 0.17 to 0.18 µm, optimized for AiryScan resolution, achieving a spatial resolution of 24 pixels/µm.

Following initial imaging, cells were processed for correlative light and electron microscopy (CLEM) according to modified versions of our own established imaging pipelines (^11,23,24^).

For serial block-face scanning electron microscopy (SEM), samples were imaged using the Gatan 3View system (Gatan) integrated with an FEI Quanta 250 FEG SEM (FEI Company). Two nested regions of interest (ROIs) were defined: **ROI_1** (full cell overview) was captured at a resolution of 4096 x 4096 pixels with a magnification of 3.6K, while **ROI_2** (chromosome region) was imaged at a higher resolution of 8192 x 8192 pixels and a magnification of 5.4K. Imaging was performed under a chamber pressure of 70 Pa and an accelerating voltage of 4 kV. Serial sections were acquired at a slice thickness of 80 nm to generate high-resolution 3D reconstructions. Lossless acquisition of >1000 sections per oocyte was performed over a continuous collection period of 72 hours.

### Analysis of volume data using Amira

#### Light microscopy Data

Light microscopy (LM) images were initially processed in ImageJ to convert raw image files into TIFF format for subsequent analysis. The processed TIFF images were imported into AMIRA software (Thermo Fisher Scientific) for detailed 3D analysis. Auto-thresholding was applied to identify chromosomes based on pixel contrast, enhancing the segmentation accuracy.

Chromosome segmentation was performed using the labelling and separate object modules within AMIRA, employing 3D interpretation parameters with a 26-neighborhood voxel connectivity and a 4-marker extension setting. Manual segmentation adjustments were made using the brush tool when necessary to refine chromosome boundaries.

Bounding boxes were applied to both the entire chromosome complement and individual chromosomes to define the area of coverage and determine centroids. This method enabled the quantification of chromosomal movement between time points, as well as calculations of total displacement and spatial dispersal of chromosomes. Additionally, the directionality of chromosome displacement and subsequent polar body extrusion were analysed using centroid data and trajectory assessments.

Statistical comparisons between control and Ki67 KD groups were performed using unpaired t-tests or Mann-Whitney U tests where appropriate, following tests for normality (Shapiro-Wilk).

#### Electron Microscopy Data

Raw electron microscopy (EM) data files in DM4 format were processed to enhance visualization and prepare them for 3D analysis. The raw DM4 files were binned at a 2 x 2 x 1 (x, y, z) ratio to optimize image resolution while maintaining data integrity. Contrast was uniformly enhanced by 0.35% across all images to improve grayscale differentiation. The adjusted DM4 files were then exported as TIFFs for analysis in AMIRA (Thermo Fisher Scientific).

In AMIRA, images were aligned to correct for any positional shifts, ensuring accurate 3D reconstruction. Auto-thresholding was applied to differentiate chromosomes based on grayscale variations. To create accurate chromosome models, the highlighted regions were manually refined to remove any excess selections.

The segmented chromosome complement was further processed using AMIRA’s labelling and separate object modules with 3D interpretation settings, including a 26-neighborhood voxel connectivity and a 4-marker extension. Where necessary, additional manual refinements were made using the brush tool to achieve precise segmentation of individual chromosomes.

### Direct comparison of EM and LM data – important technical note

Clear discrepancies were apparent between LM and EM data, but on only 3D and not 2D measurements. For example, surface area is significantly underestimated (Figure 4Hi) (LM = 35.8µm^2^+/-9.85, EM = 58.258µm^2^+/-19.5, P = <0.001) and volume is significantly over estimated (Figure 4Hii) (LM = 9.26µm^3^+/-3.61, EM = 6.81µm^3^+/-2.3, P = <0.03). Previous work of mitotic chromosomes revealed that this discrepancy was largely result of comparatively poor z resolution with volume LM data^11^, but also the likely over-smoothing of LM data for volume representation. The lack of difference in length and width measurements, acquired from an XY plane, supports this and minimises the contribution of any potential EM processing artefacts.

Cylindrical tolerance: Tolerance score was measured as the relative erosion of the chromosome area in vertical cross-section from a minimal bounding box using ImageJ. 2D images of chromosomes were thresholded and rotated to align the telomeric axis with the image Y axis.

Using ROI Manager, area of each thresholded chromosome and a corresponding minimal bounding box were computed. Tolerance ‘t’ was calculated as *t* = ^(*a_Bbox_* − *a_c_*)^⁄_*a_Bbox_*_, where *a_Bbox_* is the area of the bounding box and *a_c_* is the area of the chromosome cross section. For mitotic chromosomes, diameter measurements from Cisneros-Soberanis et al., 2024 were used to compute chromosome cross-section area.

### Human Oocyte Methods

#### Donation of human oocytes and embryos to research

This research was conducted as a part of a study approved by the NHS Research Ethics Committee (Indicators of Oocyte and Embryo Development, 04/Q2802/26) and under the Human Fertilisation and Embryology research licence (HFEA; R0155; Indicators of Oocyte and Embryo Development). Human oocytes deselected during *in vitro* fertilisation (IVF) or intracytoplasmic sperm injection (ICSI) cycles and therefore unsuitable for use in treatment were donated by patients at the Centre for Reproductive Medicine (CRM), University Hospitals Coventry and Warwickshire (UHCW) NHS Trust as described in Currie et al. (2022). Briefly, Informed consent was obtained voluntarily from all patients donating material for research. Clinical embryologists verified the consent forms and evaluated patients’ oocytes, deciding which samples qualified as research material. Samples include donations from patient 3696, aged 37 (Figure 1A, B) and patient 3695, aged 32, (Figure 1C).

**Figure 1:**
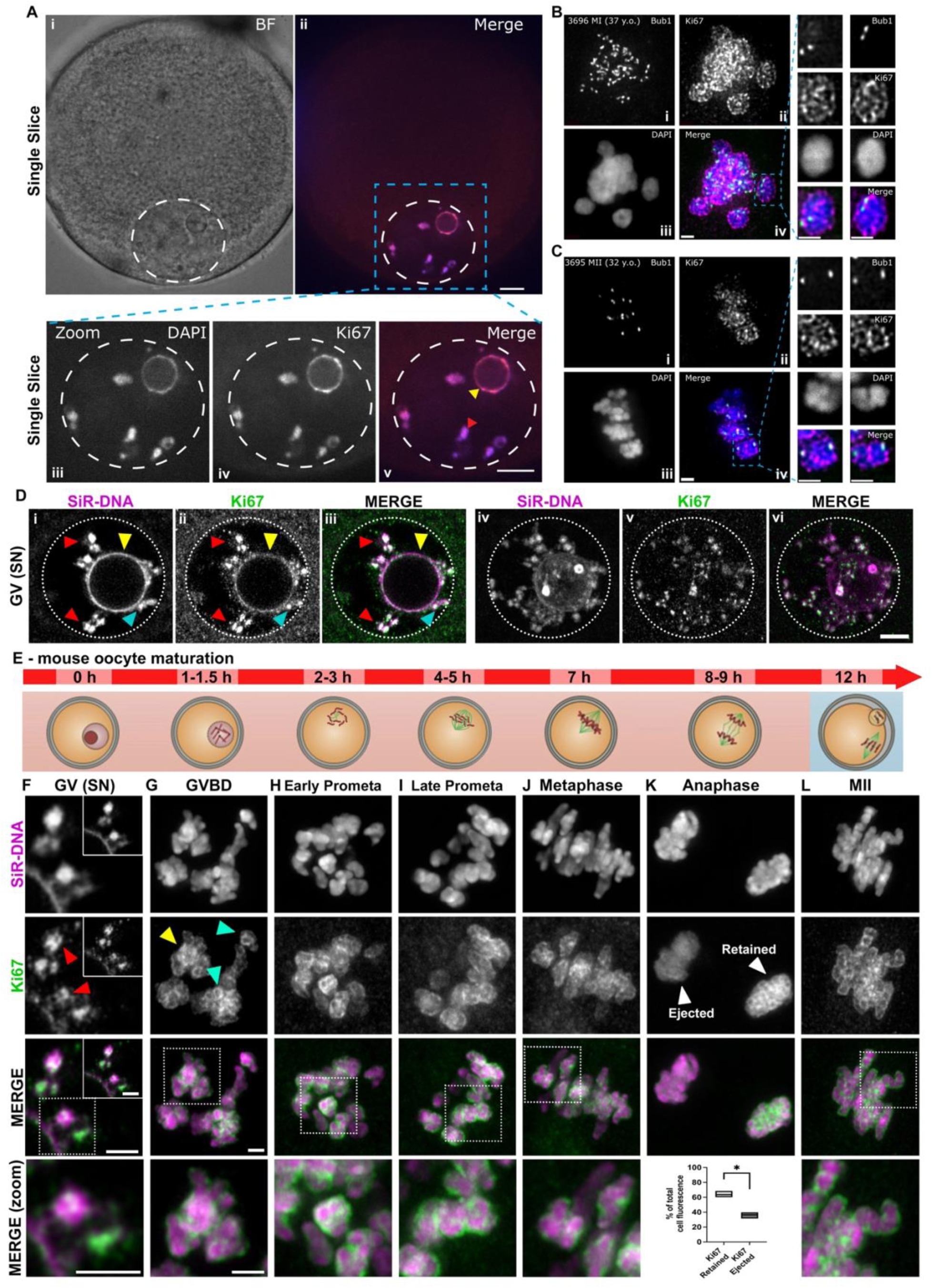
Ki67 distribution survey during meiotic progression reveals distinct differences between mitosis and female meiosis. **A**) A fixed human GV stage oocyte with a surrounded nucleolus (SN) configuration imaged using spinning disc confocal microscopy. Single slices shown. Images show overviews of brightfield (i) and merge of DAPI and anti-Ki67 (ii). Dashed circle denotes nucleus. Zoom images show DAPI (iii), anti-Ki67 (iv) and a merge (v). Arrows indicate partially condensed chromosomes (red) and nucleolar rim (yellow). Scale bar = 10μm. **B & C**) Fixed human oocytes stained for Bub1 (green, i) and Ki67 (magenta, ii), with DAPI for DNA marker (blue, iii), merge of all channels (iv). Side panels show the zoomed in area (dashed box) and examples of two sets of bivalent kinetochores. **B** shows representative Metaphase I (MI) oocyte with visible bivalents. **C** shows Metaphase II (MII) oocyte with sister chromatids organised on a metaphase plate. Scale bar = 2μm. **D**) Mouse GV stage SN oocyte fixed and probed using anti-Ki67 (green, ii, v) antibody. DNA was counter-stained with SiR-DNA (magenta, i, iv). Merge of both channels (iii, vi). Di-iii) single slice. Div-vi) Max projection. Arrows point to partially condensed chromosomes (red), nucleolar rim (yellow) or nucleolar-to-nuclear tethers (cyan). **E)** Schematic describing the staggered release/fix time-course for mouse oocytes to progress to the desired meiotic stages. **F-L)** Mouse oocytes released and fixed according to **E**. Mouse oocytes were processed for imaging as with **D**. Panels show representative projections of oocytes stained with SiR-DNA (magenta) and probed for Ki67 (green). Bottom panel shows merged zoom of the dotted area in the panel immediately above. Arrows point to Ki67 that is potentially enveloping (red), incomplete enveloping (yellow) and complete enveloping (cyan). Lower panel in **K** shows fluorescence intensity quantifications during anaphase of Ki67 from chromosome complements as they are either being ejected or retained, normalised to SiR-DNA. *N*=3^oocytes^. Bars = E-F – 2µm.

### Oocyte processing and transport

Human oocytes were cultured in Gx-TL media (Vitrolife). Zona pellucida was removed by washing with acid Tyrode’s solution (Sigma) before moving to pre-warmed, pre-gassed dishes with culture media drops overlaid with oil. All egg handling and was performed on a heated stage pre-warmed to 37°C. After processing, oocytes were placed in a humidified incubator with 5% CO_2_ to recover and then transported about 14km from the CRM to Warwick Medical School (WMS) in sealed culture dishes, in a portable incubator (K-systems) pre-warmed to 37°C.

### Oocyte fixation

Upon arrival, oocytes were fixed in formaldehyde for immunofluorescent staining as described in Mihalas et al. (2023). Briefly, oocytes were washed in 37°C PHEM buffer (60 mM PIPES, 25 mM HEPES, 10 mM EGTA, 4 mM MgSO4.7H2O; pH 6.9) with 0.25% Triton X-100 then incubated in room temperature 3.7% formaldehyde diluted in PHEM for 30 min and in PBB (0.5% BSA in PBS) for 5 min. Then oocytes were permeabilised with 0.25% Triton X-100 in PBS for 15 min and blocked in 3% BSA in PBS with 0.05% Tween-20 at 4°C overnight.

### Oocyte immunofluorescence

Fixed oocytes were incubated for 1hr in 25μl droplets of blocking solution (see above) at 37°C containing primary antibodies: mouse anti-Bub1 [2F9] (1:200, Abcam ab54893), rabbit anti-Ki67 (1:100), and human CREST (1:50, Antibodies Incorporated) overlaid with oil, followed by 15 min incubation in washing solution (PBB with 0.05% Tween-20) at room temperature. Oocytes were placed in blocking solution containing secondary antibodies: Alexa Fluor^TM^ goat anti-mouse 488, anti-rabbit 594 and anti-human 647 IgG (H+L; Invitrogen, 1:200 dilution), and DAPI (1:500) at 37°C for 1hr, and then washed again for 15 min. Oocytes were placed in 5μl drops of PBS in No. 0 coverglass bottom dishes (MatTek), overlaid with oil and stored at 4°C until imaging.

### Fixed oocyte imaging

Oocytes were imaged using Marianas spinning disk confocal microscope (3i, Intelligent Imaging Innovations) equipped with 2× Photometrics 95B Prime sCMOS cameras and controlled with Slidebook software (3i, Intelligent Imaging Innovations). Temperature was maintained at 37°C by stage-top incubator (Okolab). GV oocytes were imaged using a 63x oil immersion objective (Zeiss, 1.40 NA, Pln Apo) with brightfield, 405 (100ms exposure time, 20% laser power) and 561nm (500ms exposure time, 25% laser power) wavelengths. MI and MII oocytes were imaged under the 488, 561, 640 and 405nm wavelengths. Image stacks (200nm spacing) of area of interest (meiotic spindle) were acquired using a 100x oil immersion objective (Zeiss, 1.46 NA, alphaPlnApo) with 500ms exposure time and 20-25% laser power for 488, 561 and 405m wavelengths and 100ms exposure time and 10% laser power for 647nm.

### Image processing and analysis

Image stacks obtained using 100x objective were deconvolved in Huygens X11 software (Scientific Volume Imaging B.V.).

## RESULTS

### The first complete survey of Ki67 and CP distribution reveals key differences between oocytes and somatic cells

The presence of a CP in mammalian oocytes, and its distribution throughout female meiosis remains uncharacterised. We sought to comprehensively map the presence and localisation of the CP, using antibodies that recognise Ki67 (the organiser of the mitotic chromosome periphery), during key stages of human oocyte maturation (Figure 1A-C, Supp Figure 1A). In germinal vesicle (GV) stage ‘surround-nucleolus’ (SN) oocytes - one of the two main GV oocyte states - Ki67 co-localized with both the compact chromatin encircling the nucleolus and partially condensed prophase chromosomes (Figure 1Ai-v, yellow and red arrows respectively). Ki67 also co-localized with chromocentres that extend from the nucleolar rim and terminate at the nuclear membrane (Supp Figure 1Av, blue arrow). It is unclear why these centres are excluded from the dense heterochromatin that surrounds the nucleolus, but this is thought to support the unique epigenetic landscape in GV stage oocytes^25^.

The next assessment was post-nucleolar/nuclear disassembly - during MI (Figure 1B) and MII (Figure 1C). We report the first evidence of a CP neatly decorating the surfaces of condensed human meiotic chromosomes.

Next, we extended the survey to include mouse oocytes, reasoning that their comparatively better availability would allow wider coverage of maturation stages. Indeed, whilst confocal imaging of Ki67 in mouse SN GV-arrested oocytes (Figure 1D) mirrored that of human, there was surprising variability of Ki67 distribution between SN and ‘non-surrounded-nucleolus’ (NSN) GV oocytes (Compare Figure 1D and Supp Figure 1B). A defining feature of SN oocytes is a tight chromatin ring surrounding the nucleolus (Figure 1Aiii, Di), which is absent in NSN oocytes. Interestingly, Ki67 was also absent (Supp Figure 1B) – quantified using line-scans (Supp Figure 1Ci,ii). SN and NSN are discrete morphologies in oocytes that describes the maturity of the oocyte towards the end of the prophase I arrest period, a key distinction as SN/NSN status is heavily linked with oocyte viability. Importantly, the localisation of Ki67 in both SN and NSN oocytes, excluded from the nucleolus, is in stark contrast with somatic cells, where almost all Ki67 is confined within the nucleolus, and indeed, sequestered away from chromatin, until nuclear envelope and nucleolar disassembly^26^.

We continued to survey the CP during key stages of mouse oocyte maturation (Figure 1E). Although partial ‘enveloping’ was noted in some GV oocytes (Figure 1F, red arrows), recruitment of Ki67 to chromosomes was most pronounced from GVBD, where it fully enveloped some chromosomes (Figure 1G, cyan arrows) but still only partially coated (i.e. a continuous rim cannot be viewed regardless of the angle or z position) others (Figure 1G, yellow arrow). As oocytes progressed from early prometaphase (Figure 1H), through late prometaphase (Figure 1I) and to metaphase (Figure 1J), Ki67 remained tightly associated with the surface of all condensed meiotic chromosomes – closely resembling their somatic cell mitotic CP counterpart. However, a remarkable variation, potentially unique to meiosis, was observed during anaphase I, where Ki67 was unequally distributed between the chromosomes retained in the oocyte, versus those to be ejected into the polar body (PB) (Figure 1K). Ki67 fluorescence intensity measurements revealed that 63.6% (±4.3) of Ki67 remained on chromosomes segregating towards the oocyte, while 36.4% (±4.3) was found on chromosomes segregating towards the PB (Figure 1K, lower panel graph. P = >0.05). Furthermore, Ki67 no longer fully enveloped the PB-facing chromosomes, instead decorating chromosomes more diffusely. Notably, no significant difference was observed between the intensity of DNA between retained and PB-facing chromosome sets (Supp Figure 1D). Finally, Ki67 was still recruited neatly to the surface of MII oocyte chromosomes (Figure 1L, and also in human Figure 1C).

### Depletion of Ki67 reveals a novel role for the CP in meiotic progression

We next sought to assess the function of the meiotic CP by depleting Ki67 and following chromosome segregation by live imaging mouse oocytes (Figure 2, Supp Video 1-4). Using a morpholino oligo we depleted Ki67 by 66% relative to typical wildtype (Supp Figure 2A-C). Ki67 depletion had a substantial impact on the oocytes ability to progress to anaphase, as only 62% of oocytes produced a polar body, compared to 89% of controls. Many depleted oocytes stalled, displaying alignment, congression and “clumping” defects (Figure 2A, B, Ci, Supp Figure 2D). Of the depleted oocytes that did progress to anaphase, 66% of these oocytes managed an apparently faithful segregation, compared to 75% of controls. In addition, they did so with normal timings – as mean GVBD duration to polar body formation showed no significant difference between control (∼533±99.2 minutes) and Ki67 depleted oocytes (∼512±97.1 minutes) (Figure 2Cii). Although several of the knockdowns exhibited gross mis-segregation errors (Figure 2A far right).

**Figure 2:**
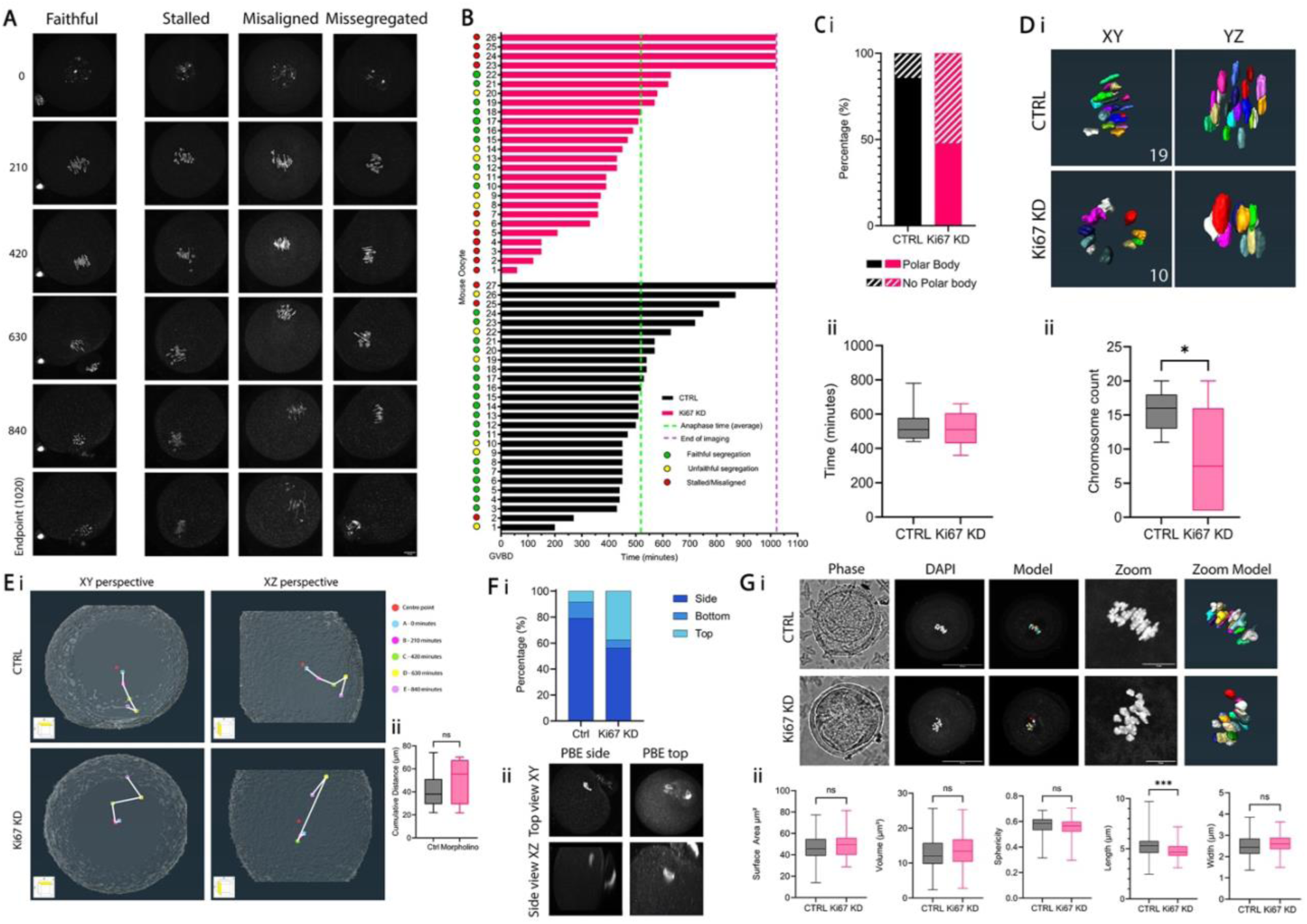
Ki67 knockdown causes errors in meiotic progression. **A)** Representative “stills” lifted from time-lapse imaging sequences of oocytes, in control or following injection of Ki67-targeting morpholinos and staining with SiR-DNA. Meiotic progress was scored as one of four criteria: faithful (successful), stalled, misaligned and mis-segregated (in this example a PB was formed but the full set of chromosomes were ejected) (see methods for scoring criteria). Scale bar = 10 µm. **B)** Bar chart showing quantification of duration (minutes) from germinal vesicle breakdown (GVBD) to anaphase onset, comparing control (black) and Ki67 KD (pink) oocytes. Oocyte division outcomes were classified as faithful (green), unfaithful (yellow). Stalled/misaligned cells (red) did not divide and either failed to progress to polar body (PB) formation or underwent apparent cell ‘death’. *N*=27^CTRLoocytes^ *N*=26^Ki67KDNoocytes^. **C)** Analysis of meiotic progression frequency from oocytes in panel B, showing the percentage of oocytes that achieved PB formation in control versus Ki67 KD groups (Ci). Cii) Bar chart showing mean duration from GVBD to anaphase onset for all “faithful” oocytes. (*P > 0.05*). **D)** Stills for each oocyte were lifted from time-lapse videos at prometaphase. 3D models generated using AMIRA software were used for quantification of individual chromosomes present in each oocyte **(i)**. **(ii)** Plot depicting chromosome counts in control and Ki67 KD cells, with significantly more individual chromosomes visible in control cells (*P = 0.039*). **(E)** Representative models, from two perspectives, for movement of chromosomes tracked throughout the cell from GVBD onwards. Dots represent the centroid of the chromosome complement and colours depict time points. (Eii) Bar chart of mean distance travelled for chromosome complements. **(F) (i)** Directionality of polar body extrusion (PBE) was assessed in control and Ki67 KD oocytes, scored as side, bottom or top. **Fii)** Representative images illustrating PBE sites. **(G)** Super-resolution microscopy imaging and analysis of control and knock down oocytes released to metaphase. **Gi)** Representative images of fixed mouse oocytes stained with DAPI and imaged using AiryScan LSM900, along with corresponding segmented chromosome complement models (far right panels). **Gii)** Geometric analysis of segmentation data, including total surface area, total volume, mean chromosome length, mean chromosome width - comparing control (*N*=7^oocytes^) and Ki67 KD (*N*=7^oocytes^) cells. Data are presented as mean ± SD unless otherwise stated. Bar = 10 µm.

Due to the substantial disruption to meiotic progression following Ki67 knockdown, we considered whether a mechanism linked to chromosome dynamics, structure, or individualisation had been impacted by Ki67 depletion. 3D digital reconstruction of chromosomes from our time-lapse data allowed us to assess chromosome individualisation (Figure 2D) and dynamics (Figure 2E-F). Modelling of mid-prometaphase chromosomes (Figure 2Di) - the period of cell division we quantified control chromosomes to be maximally dispersed - revealed that far fewer chromosomes were observable in depleted oocytes (8.5 ± 7.5) versus those of controls (15.8 ± 3.0) (Figure 2Dii), indicating issues with maintenance of chromosome individualisation. This had little impact on overall chromosome dynamics as tracking revealed no significant change in sum distance travelled between chromosomes of control (41.18µm ±15.84) or Ki67 depleted (50.75µm ±20.80) oocytes (Figure 2Ei-ii). However, we did note a highly unusual alteration in direction of polar body formation and chromosome ejection (Figure 2F). In control oocytes chromosomes were most frequently ejected from the side of the oocyte (80%) as would be typical for cultured oocytes in this setting. In contrast, in Ki67 depleted oocytes only 57% side ejected, and instead, a large portion (38%) ejected from the top of the oocyte (Figure 2Fi-ii). This is contrary to what we understand about the forces that govern the positioning of the spindle in-vitro, and it is difficult to appreciate how an upwards trajectory of chromosomes might be energetically favoured.

Using super-resolution imaging of fixed samples, we next determined whether Ki67 depletion impacted chromosome morphology (Figure 2G). No difference was found between control and knockdown chromosomes for chromosome width (2.5 vs 2.6µm), volume (12.7µm^3^ vs 13.6µm^3^), or surface area (46.9µm^2^ vs 49.56µm^2^). This data, in particular the volume, strongly suggests that although chromosomes are closer together (less individualised), they are not hyper-condensing. Interestingly however, a significant difference was determined in mean chromosome length, with knockdown chromosomes being slightly shorter (5.1µm vs 4.4µm), suggesting some aspects of chromosome morphology are disrupted. During characterisations of the mitotic CP, we found it essential to perform an ultra-structural analysis^9,11^. Therefore, we moved to 3D correlative light electron microscopy (3DCLEM), to better determine the molecular mechanism underpinning meiotic Ki67 function.

### Ultra-structural analysis of Ki67 depleted oocytes reveals multiple key features

We previously validated 3DCLEM as a powerful multimodal imaging tool for investigating mitotic chromosome architecture and morphology^11,24^ and adapted this here as a platform for studying oocytes (Figure 3, Supp Videos 5,6). This is non-trivial as, unlike mitotic chromosomes, mouse oocyte chromosomes only account for ∼0.07% of the entire oocyte volume (calculated from our own 3DCLEM data – sum chromosomes volume = 136µm^3^, oocyte volume = 187,401,136µm^3^). Therefore, we used fluorescence data (SiR-DNA) to guide us, in 3D Euclidian space, to areas of interest within the oocyte (Figure 3Ai-iii,Bi-iii), i.e the regions containing chromosomes, and acquire high resolution volume data (Figure 3Aiv-vi,Biv-vi). We also used the optical images as a “rare event” finder, helping us pre-select cells with a ‘moderate’ phenotype, reasoning that this would yield the most useful data.

**Figure 3:**
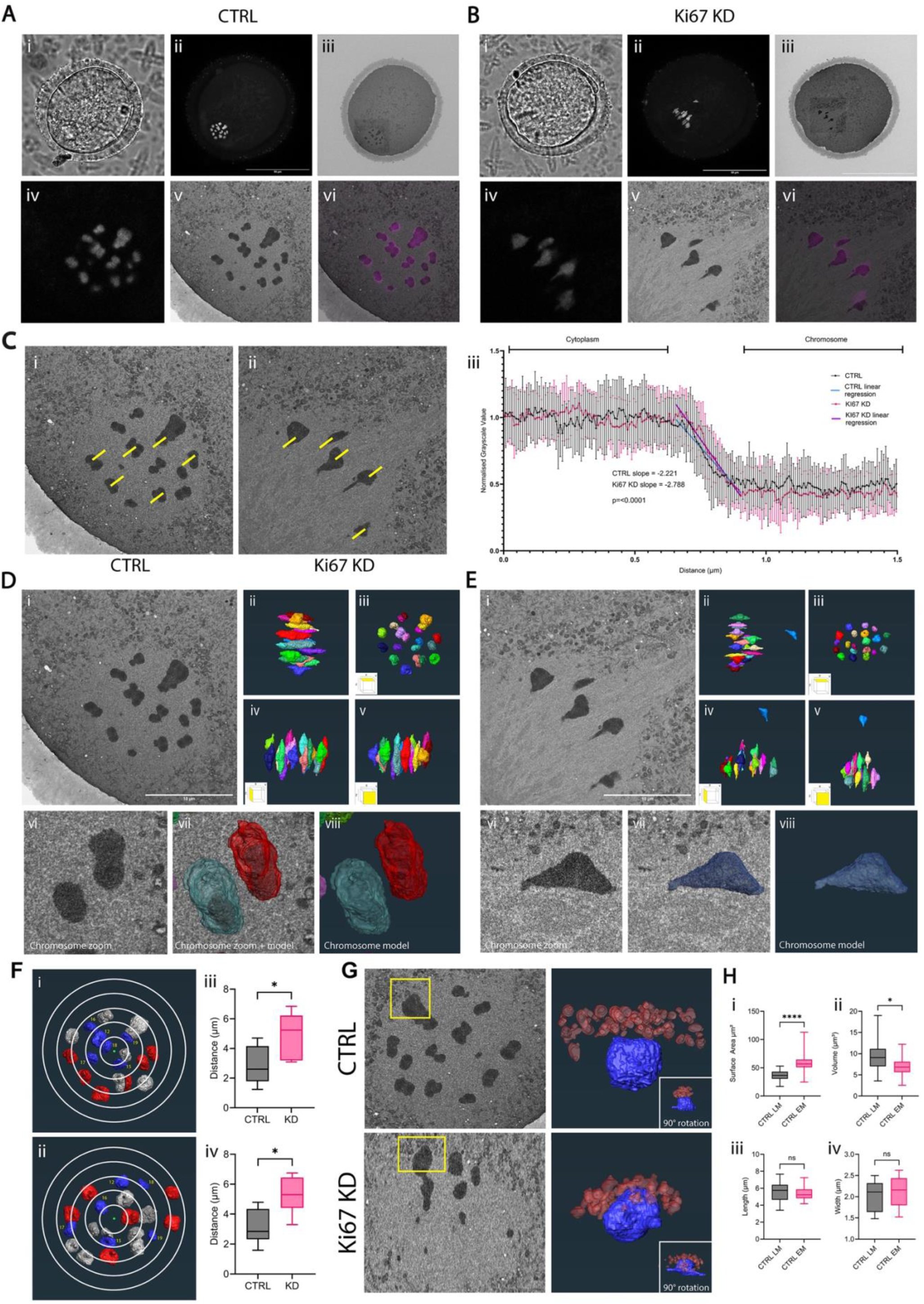
Correlative Light and Electron Microscopy (CLEM) analysis of oocyte chromosomal architecture reveals multiple key findings. **A-B)** High-resolution CLEM imaging of a control oocyte (A) or Ki67 KD oocyte (B), showing whole oocyte overviews of phase contrast (i), SiR-DNA (ii) and electron microscopy (iii). High resolution of the area of the chromosomes was also acquired showing, SiR-DNA (iv), EM (v) and an LM to EM overlay to confirm registration (vi). **(C)** Comparative line scan analysis to measure pixel density of the chromosome periphery. 1.5µm line scans (yellow), originating in the chromosome body and terminating in the cytoplasm, were used to retrieve pixel density every 10nm. **(i)** representative control oocyte, **(ii)** representative Ki67 KD oocyte, **(iii)** grayscale plot with values normalised to cytoplasm measurements, slope analysis using linear regression analysis and ANCOVA, revealing a significantly steeper slope (*P < 0.0001*) in Ki67 KD cells, indicating effective chromosome periphery removal. *N*=10^chromosomes^ *N*=50^line-scans^. **(D-E)** AMIRA-generated 3D models of chromosome complements in control (D) or Ki67 KD (E) oocytes: **(i)** corresponding EM image, **(ii)** metaphase chromosome alignment, **(iii)** XY perspective, **(iv)** XZ perspective, **(v)** YZ perspective, **(vi)** magnified view of selected chromosome(s), **(vii)** overlay with AMIRA model, and **(viii)** isolated AMIRA model for detailed structural visualisation. **(F)** left panels) EM slices illustrating chromosome positioning relative to exclusion zone boundaries. Yellow boxes indicate chromosomes closest to the edge of the exclusion zone. In control oocytes, chromosomes remain confined within the boundary, while in Ki67 KD oocytes, chromosomes penetrate beyond this boundary, with partial “ensheathing” by membranous organelles. **F)** right panels) Segmentation of membranes to illustrate ensheathing. **(G)** Assessment of chromosome distribution relative to chromosome volume. Distance from chromosome centroid to centre of metaphase plate was measured for the 6 smallest chromosomes (blue) for control (i) and knockdown (ii) chromosomes, in both 2D (iii) and 3D (iv). 2um spaced concentric rings superimposed to illustrate the redistribution of large (red) and small (blue) chromosomes. Remaining chromosomes are painted grey. **(H)** Geometric analysis comparing correlative LM and EM measurements in control CLEM samples. Bar charts show mean chromosome; surface area, volume, length and width.

3DCLEM and nanometre precision 3D reconstructions of each and every mouse oocyte chromosome revealed several important features, including:

1. Depletion of Ki67 leads to a physical loss of the meiotic periphery compartment (Figure 3C). Line scan profiles that traversed the chromosome to cytoplasm interface (i.e. containing the CP) revealed a statistically significant difference between linear regression profiles of control and knockdown chromosomes (control slope = −2.221, knockdown slope = −2.788, P=<0.0001), with the latter showing a steeper pitch (Figure 3Ciii). Our interpretation of this is the physical removal of the chromosome periphery layer, supporting our Ki67 knockdown data (Supp Figure 2) and previous work performing a similar analysis of mitotic chromosomes^9^.
2. In control oocytes the smaller chromosomes tended to reside towards the centre of the metaphase plate and larger chromosomes more distal (Figure 3Fi). A positional trend consistent with other studies of both mitotic^11,27,28^ and meiotic chromosomes^29^. However, following Ki67 depletion, this size-based spatial-distribution is disrupted, with most of the smaller chromosomes now residing towards the exterior of the chromosome set (Control = 2.85µm±1.29, Ki67KD = 4.94µm±1.51, P= <0.03) (Figure Fii, iii). This trend was significant in proximity measurements performed in both 2D (Figure Fiii) and 3D (Figure Fiv), therefore taking into consideration depth of the metaphase plate. The necessity of this volume related spatial re-arrangement remains unclear, but could suggest a mass/size-based vulnerability, with smaller chromosomes more likely to succumb to abnormal “repulsion” forces exerted by the CP. This finding appeared to trend with an additional, observation of “encroaching of membranes” (Figure 3G).
3. A comparison of volume EM data from control and Ki67 depleted oocytes revealed the unexpected finding that membranes and membranous organelles (most notably mitochondria), typically excluded from chromosome/spindle regions, tended to partially envelope some of the Ki67 depleted chromosomes (Figure 3G), most notably on chromosomes towards the exterior of the chromosome complement. This suggests the CP might also function to “repel” objects other than solely chromosomes.
4. We next exploited the nanometre precision of our 3DCLEM data to create a digital karyotype – plating out models of each chromosome, ranked by volume and DNA content (Supp Figure 3). In contrast with their mitotic counterparts^24^, meiotic oocyte chromosomes displayed striking variability in morphology (in particular the clearly misaligned chromosome in the Ki67 KD cell). We scored cylindrical tolerance (see methods) for chromosomes where a perfect cylinder has a tolerance of 0, finding mitotic chromosomes to have a score of 0.18 ± 0.03 while meiotic chromosomes were scored as 0.4 ± 0.05. Thus, whilst mitotic chromosomes have frequently been modelled as cylinders (with round capped-ends), the same coarse analysis cannot be applied to that of meiotic oocyte chromosomes. To this end we also highlight an important technical note – that the retrieval of chromosome geometry measurements, using light microscopy data only, should be performed with caution. A direct comparison of super-resolution light microscopy data with volume electron microscopy data, of the same control oocyte, revealed stark inaccuracies with LM measurements (Figure 4H), see methods for details.

**Figure 4:**
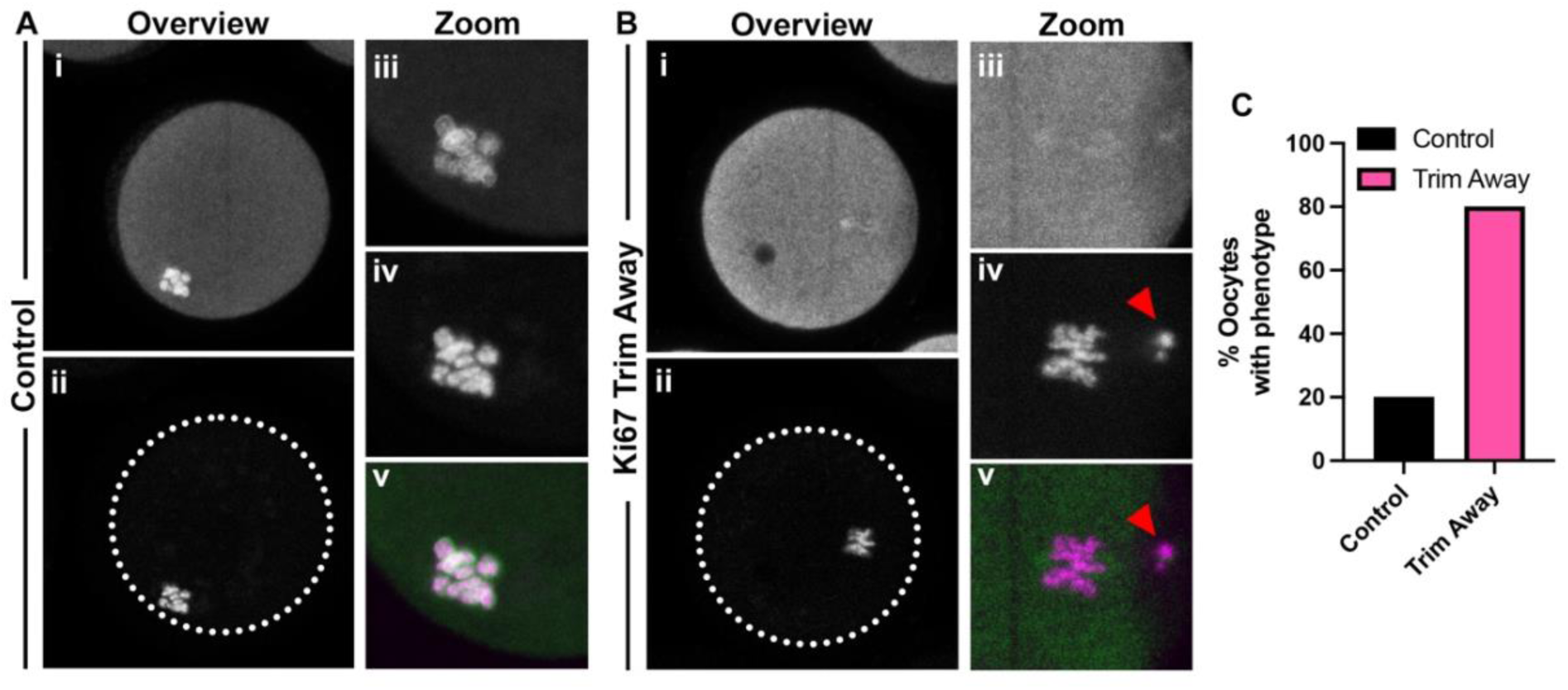
Acute depletion of Ki67 following GVBD causes meiotic defects. **A-B)** Oocytes were treated using “Trim-away” as controls (A) or Ki67 depleted (B). Oocytes were released for 16 hours, fixed and probed for Ki67 (green, i, iii, v) and counter stained with SiR-DNA (magenta, ii, iv, v) before confocal imaging. Representative images of control (A) or Ki67 depleted oocytes (B) are shown. Oocyte overviews (i-ii) and zooms of chromosomes (iii-v) are shown. Arrows point to misaligned or lagging chromosomes. C) Quantification of images in A and B. Oocytes were scored for defects, including, misaligned chromosomes and lagging chromosomes. N=5^cnt^ ^oocytes^ N=5^KD^ ^oocytes^.

### Acute Depletion Confirms Novel Ki67 Function, Explicitly During Meiotic Progression

In somatic cells, multiple cell-cycle stage dependant functions have been reported for Ki67 in interphase (i.e. pre-nuclear envelope breakdown). Therefore, we next sought to determine whether our new discoveries in meiosis were attributed to a role for Ki67 before or after GVBD – i.e. a function within the oocyte nucleus or instead at the surface of fully condensed chromosomes. We used TrimAway^30^ for acute Ki67 removal specifically following nuclear envelope breakdown, and assessed for potential abnormal phenotypes using fixed imaging microscopy (Figure 4A-C). Remarkably, almost all oocytes (80%) that underwent acute depletion of Ki67, displayed an abnormal meiotic phenotype, such as: chromosomes coalescing or misalignment/lagging chromosomes (Figure 4B, C). In contrast, only 20% of control oocytes showed abnormalities (Figure 4A, C). This suggests that Ki67’s key function in oocytes is predominantly performed as a component of the meiotic chromosome periphery.

This study has provided the most comprehensive survey of Ki67 distribution in female meiosis to date, and also the first analysis of Ki67/CP function(s) in oocytes. We reveal novel features of Ki67 localisation, relating to both pre and post GVBD in addition to previously unknown roles in faithful meiotic chromosome segregation, in the establishment of an aligned metaphase plate and the coordinated segregation of chromosomes.

## DISCUSSION

### Surveying the Distribution of Ki67 and the CP During Female Meiosis

Here we provide the first evidence for a chromosome periphery compartment in human and mouse meiotic oocyte chromosomes – and demonstrate that this is critical for normal chromosome segregation.

We begin by demonstrating a clear difference in Ki67 distribution in two very different, but readily observable GV oocyte states. At the GV stage, alongside transcriptional silencing, chromatin morphology alters dramatically. In the actively transcribing and rapidly growing oocyte, chromatin is uncondensed, dispersed across the GV, often described as ‘non-surrounded nucleolus’ (NSN). Then, as the oocyte is prepared for ovulation, chromatin is reorganised. The now heterochromatin, condenses, enveloping the nucleolus with morphology described as ‘surrounded nucleolus’ (SN)^31^. We observed Ki67 to be strongly associated with chromatin in both SN and NSN oocyte states. The significance of this is unclear, but is likely important because while oocytes with either chromatin arrangement are able to resume meiosis, they do not have comparable developmental potential - embryos derived from NSN oocytes often halting development at the two-cell stage, while embryos from SN oocytes are capable of developing to the blastocyst stage and full term^31^. Of additional interest, in somatic cells almost all Ki67 is held *inside* the nucleolus, remaining there until nucleolar and nuclear envelope breakdown at mitotic entry, at which point it is one of, if not the first, of a group of 50+ nucleolar proteins to re-localise to the surface of condensing chromosomes^26,32^. In contrast, Ki67 in GV oocytes resides *outside* of the nucleolus, indicating a basis for alternative pre division functions of Ki67, between mitosis and meiosis but also alternative mechanisms of initial CP-to-chromatin engagement.

Following GVBD, Ki67 distribution through to MII was largely similar to mitosis, but for one key distinction – meiotic Ki67 was, surprisingly, unequally distributed between chromosome complements during anaphase – with significantly more Ki67 retained than is expelled in the polar body. This is particularly curious given that during mitosis, Ki67 and the CP act as a scaffold for nucleolar proteins to “piggy back”, in equal abundance, into the new daughter cells^9^. Clearly, there is no identical daughter cell production in female meiosis, however there is fertilisation at MII by mature sperm, which have undetectable levels of Ki67^22^. Thus, a meiotic refinement of the division process could be pre-empting the need for additional Ki67, and potentially other nucleolar proteins, not supplied by sperm.

### A Novel Role for Ki67 and the CP in Meiotic Progression

We went on to demonstrate a critical role for Ki67 during meiotic progression, where Ki67 depletion resulted in a significant drop in meiotic progression and inhibition of faithful chromosome alignment and segregation. 3D reconstructions of the chromosomes revealed that fewer chromosomes could be counted following Ki67 depletion but little difference between overall chromosome volume (i.e. no evidence of hyper compaction), suggesting that removal of the CP in oocytes causes chromosome to aggregate or “clump” within a smaller area, losing the ability for “individualisation” and thus could contribute to halting meiotic progression. Interestingly, the same chromosome “clumping” phenotype is, with no mechanism assigned, prevalent in human oocytes that stall on approach to fully aligning a metaphase plate^4^. Given that Ki67 abundance is so highly variable in WT oocytes even from the same laboratory mouse (Supp Figure 2A-C), it is feasible that an optimal abundance of Ki67 is critical, with deviations from this contributing to these naturally occurring clumping events in human. This is supported by recent work in rhesus monkeys reporting significant decrease of Ki67 mRNA, in stalled oocytes, relative to normal^33^.

Curiously, chromosome clumping was also present in several of our control oocytes (Figure 2B), which in some cases, actually exhibited such naturally low Ki67 levels, they were comparable with those of knockdown. This presumably relates to variable starting levels of Ki67 and would explain why some oocytes, with naturally high abundance of Ki67, might have been less impacted by depletion.

Of the Ki67 depleted oocytes that did progress to anaphase, we noted a clear change in the site of chromosome ejection. Typically, a polar body forms at the side of oocytes, however, we observed a significant shift towards ejection at the top of the oocyte. The cause of this abnormality remains unclear, but could be related to observed defects in chromosome morphology (length) or loss of individualisation. Alternatively, or perhaps additionally, a mechanism might stem from the unequal retention of Ki67 and instead link to meiotic drive and spindle asymmetry. Here, half of the spindle is more “selfish”, a phenomenon thought to confer genetic inheritance advantages and cell diversity^34,35^. This could align with our Ki67 knockdown data showing an apparent loss of spatial awareness of the chromosomes (and presumably spindle), resulting in abnormal exit position of the polar body. Moreover, the chromosomes themselves appeared to lose intra-chromosomal positional awareness, with an abnormal redistribution of smaller chromosomes towards the outside of the metaphase plate. The significance of this remains unclear but is consistent with a broader mechanism recently reported between chromosome length changes (stretching) and oocyte chromosome positioning^29^. This is particularly interesting given the preferential loss or gain, specifically of smaller chromosomes, in certain conditions. For example, preferential loss of smaller chromosomes in solid tumours^36^, but also that, in oocyte aneuploidy, the preferential gain of only chromosomes 13, 18 and 21, can result in a viable foetus.

Importantly, we were able to confirm that our novel findings were not simply the result of chronic Ki67 depletion prior to meiosis resumption, i.e. whilst a nucleus and nucleolus was still intact. We bypassed this possibility using acute depletion by Trim-Away^30^. This requires antibody microinjection into the oocyte cytoplasm, where it binds to the target protein and therefore marked for proteasomal degradation. As the antibody is too large to move through the nuclear envelope it would have no access to Ki67, until GVBD. Therefore, in these experiments the increase in defects observed for acute Ki67 depletion can only be attributed to inhibition of a role in prometaphase. This was important to determine since in mitosis, numerous interphase (nuclear/nucleolar) roles for Ki67 have been reported, included those linked to proliferation^37^.

### 3DCLEM improves our understanding of mouse oocyte chromosome ultra-structure and the role of Ki67

This allowed us to assess a variety of useful, previously unattainable, ultra-structural features, including; confirmation of physical removal of the CP, disruption to spatial distribution of chromosomes, encroachment of membranes to chromosomes, and finally, a unique insight into overall variability in meiotic chromosome morphology.

Following Ki67 depletion, we noted a striking partial enveloping of chromosomes with membranous organelles. Given that the CP has a role in mitigating chromosome-to-chromosome aggregation, it is reasonable that same function could extend to also include none chromosome objects – protecting chromosomes against coating/aggregation by membranes, which could be problematic. Indeed, in mitotic cells, a phenomena of aberrant “membrane ensheathing” has been linked to chromosome segregation defects^38^. In the context of our work, this may be a contributing factor to the abnormal ejection of meiotic chromosomes during polar body formation and could be an important novel link between Ki67 and a mechanism underpinning oocyte viability. It is tempting to speculate that Ki67 or the CP could now be explored for therapeutic benefit, indeed, Ki67 is already actively being pursued as a therapeutic target for cancer^39^. How these therapies progress and how relevant they might be in a reproductive context, remains to be seen, but is an exciting prospect. Regardless, our work and bespoke imaging pipelines have revealed compelling evidence of a new, chromosome periphery dependant pathway, that is critical for oocyte maturation, further contributing to our fundamental understanding of female meiosis.

## AKNOWLEDGEMENTS

**DGB** thanks; BBSRC David Phillips Fellowship (**DGB** and **XX**, BB/V005626/1), Royal Society (RGS/R2/202366), Academy of Medical Sciences Springboard Award (**DGB**, SBF006\1071) and Leverhulme Trust (**TM**, RPG-2021-118). **ES** is supported by a BBSRC PhD studentship. **SM** and **BW** was supported by an MRC Career Development Award [MR/T010789/1]. **HM** and **LPC** were supported by European Research Council (ERC) consolidator grant CODECHECK, under the European Union’s Horizon 2020 research and innovation programme (grant agreement 681443) and Fundação para a Ciência e a Tecnologia of Portugal (EXPL/BIA-CEL/0604/2021). **LPC** was funded by a Marie Skłodowska-Curie Actions Fellowship (746515). **IAP** is supported by a North West Cancer Research endowment (CR1166). **ADM** and **GMH** are supported by a Wellcome Collaborative Award (215625). **GMH** is supported 50% by UHCW NHS Trust.

---

**Supplementary Figure 1:**
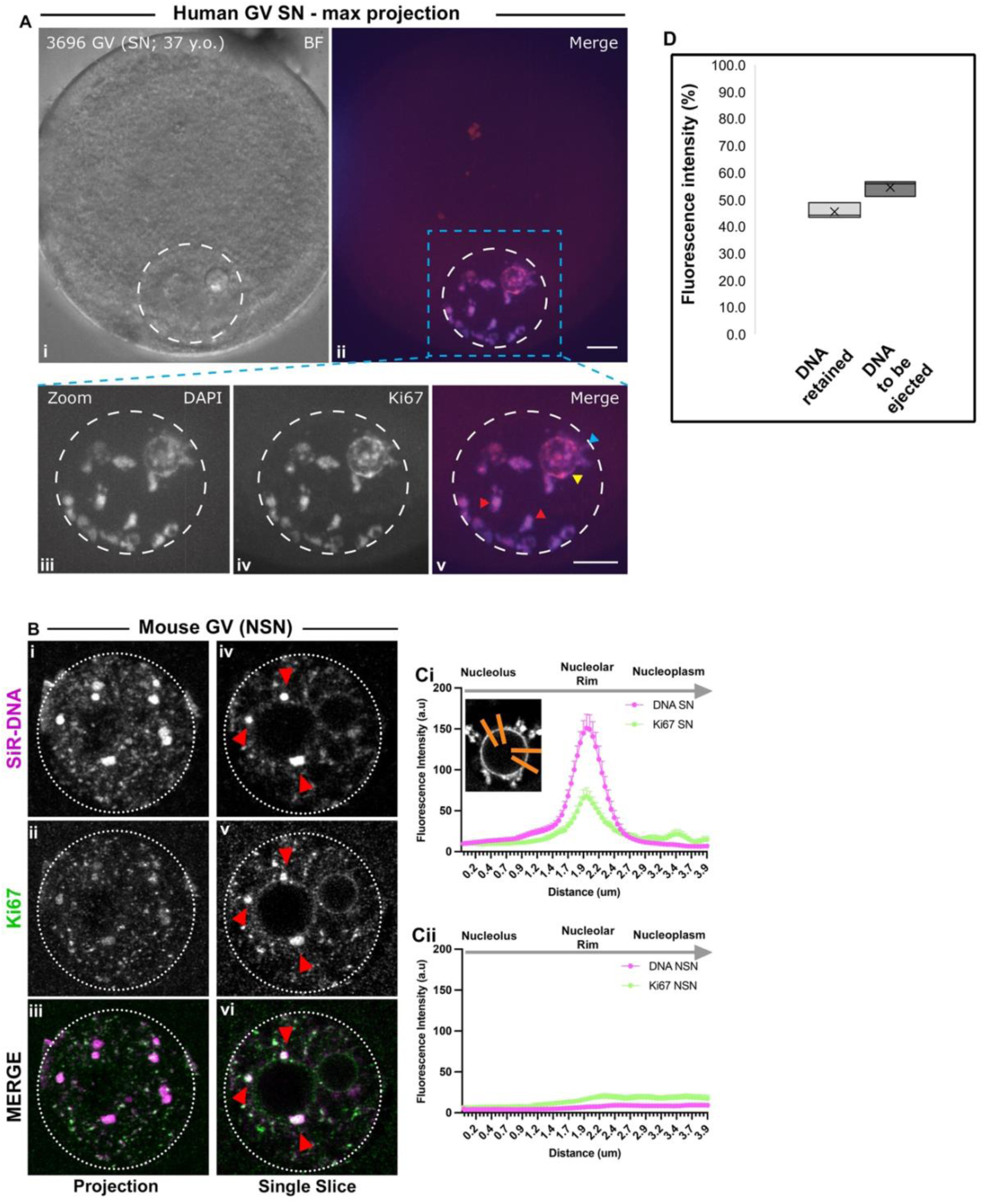
Quantification of SN and NSN oocytes. **A**) The same human oocyte as in Figure 1A, but shown here as a max projections in all channels. Arrows point to partially condensed chromosomes (red), nucleolar rim (yellow) or nucleolar-to-nuclear tethers (cyan). **B**) Mouse GV NSN oocyte fixed and probed using anti-Ki67 (green, ii, v) antibody. DNA was counter-stained with SiR-DNA (magenta, i, iv). Merge of both channels (iii, vi). Bi-iii) single slice. Biv-vi) Max projection. Arrows point to partially condensed chromosomes (red). **C)** Quantifications of SirDNA and Ki67 from images in Figure 1D for SN oocyte and Supp Figure 1B for NSN oocyte. 4µm line scans were placed (in non-occluded areas) traversing the interface of the nucleolus and into the nucleoplasm, with the nucleolar rim at the centre of the line scan (see Ci inset for example). Pixel densities were acquired every 25nm. *N*=3^oocytes^ *N*=24^line-scans^. Bar = 5µm. D) Quantifications of SiR-DNA from mouse oocytes in anaphase. Total fluorescence intensity measured shown as % split between the retained chromosome set and ‘to be’ ejected chromosome set.

**Supplementary Figure 2:**
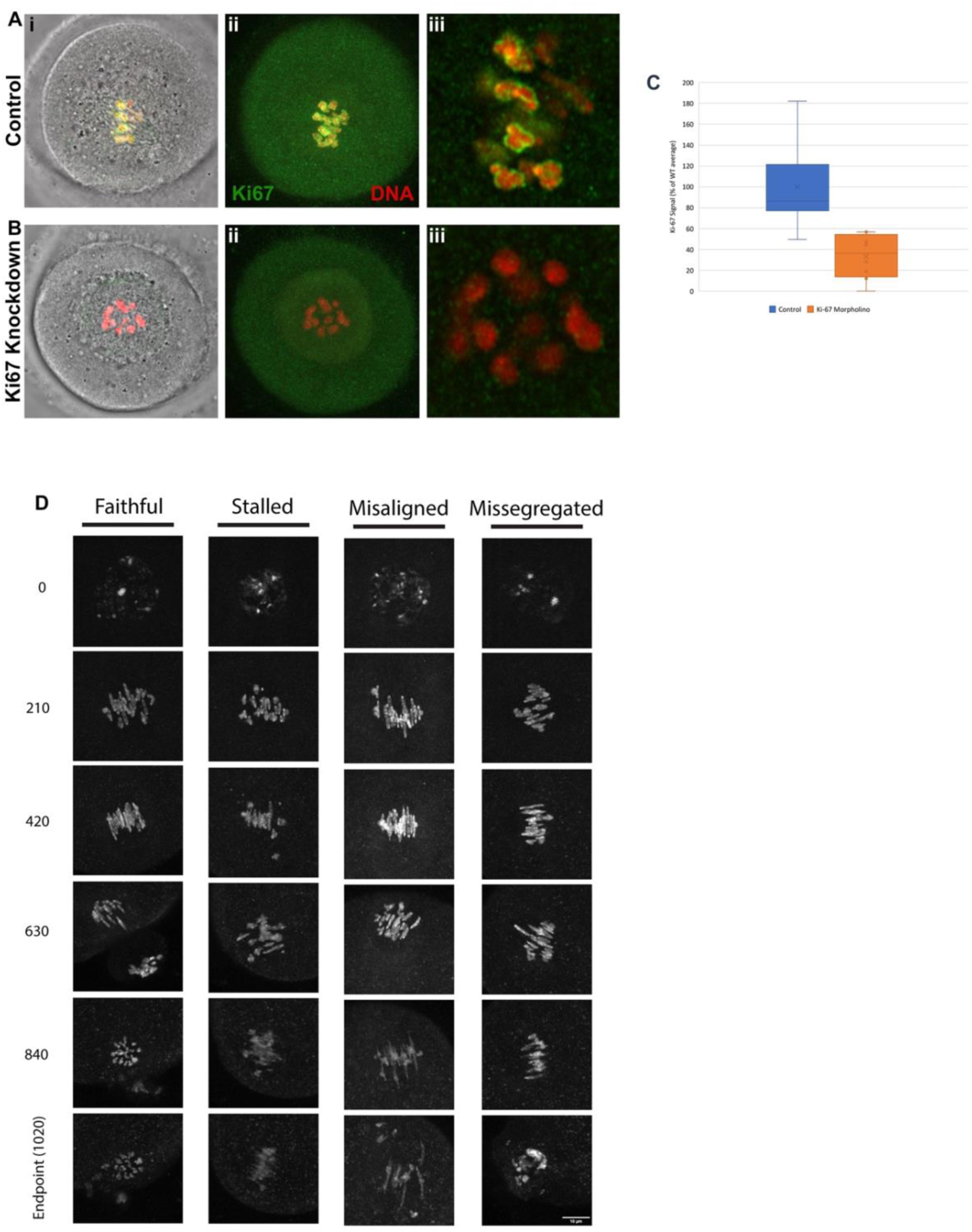
Quantification Ki67 abundance following knockdown using morpholinos. **A)** Oocytes were held in GV, and either remained as control or were injected Ki67 targeting morpholinos, prior to release for 6 hours. Oocytes were fixed and probed with anti-Ki67, stained with SiR-DNA and imaged by confocal microscopy. Representative images of control (A) and Ki67 knockdown (B) oocytes are shown. C) Quantification of Ki67 fluorescence intensity. N=5^cnt^ ^oocytes^ N=5^KD^ ^oocytes^. D) Zooms of live imaging stills, showing representative examples of faithful, stalled, misaligned and missegregated oocytes, from Figure 2A.

**Supplementary Figure 3:**
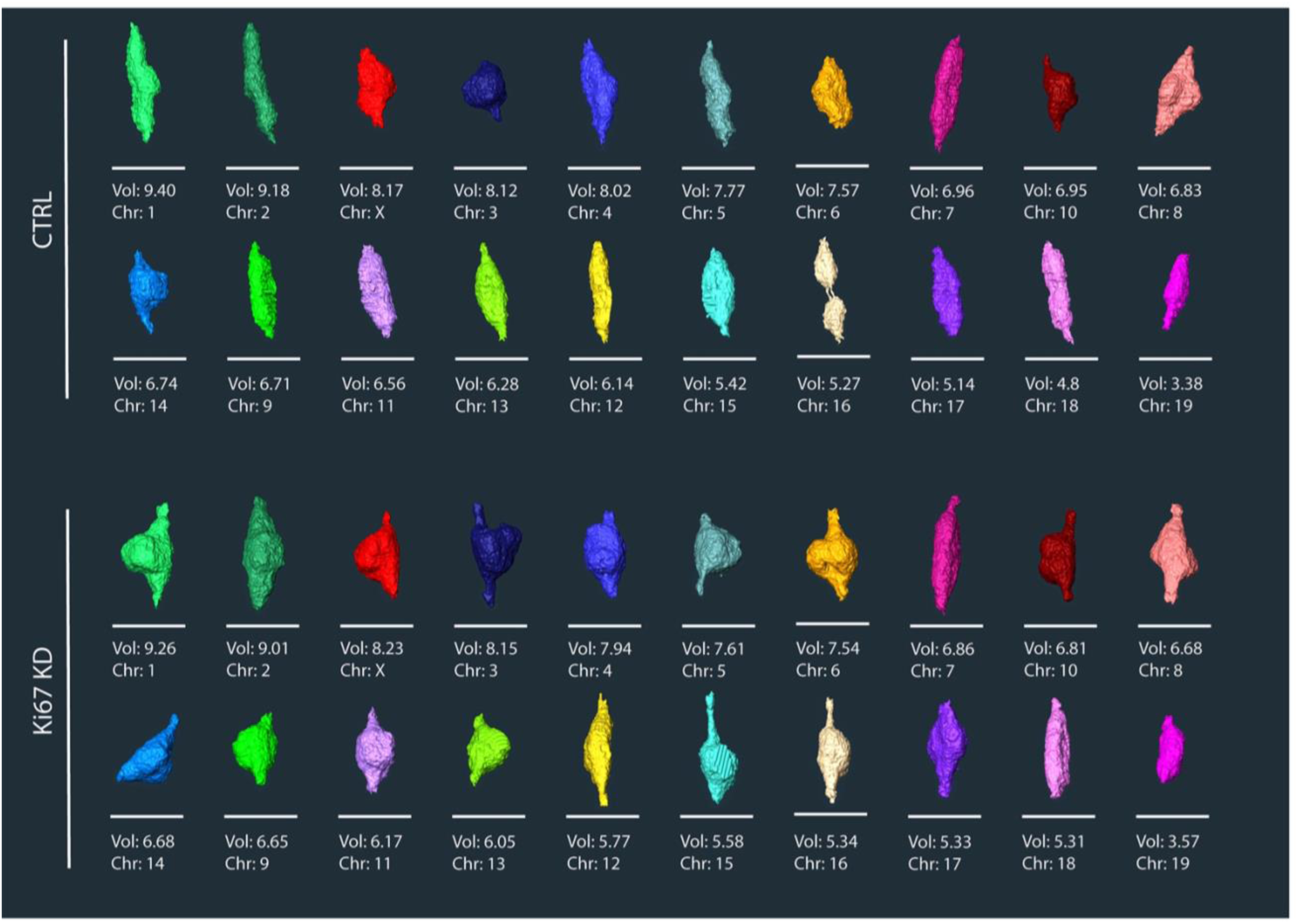
Digital 3D “karyotype” of reconstructed oocyte chromosomes. All 20 chromosomes from control oocytes (A) or Ki67 depleted oocytes (B).

## REFERENCES

1 Webster, A. & Schuh, M. Mechanisms of Aneuploidy in Human Eggs. Trends Cell Biol 27, 55–68, doi:10.1016/j.tcb.2016.09.002 (2017).

2 Hassold, T. & Hunt, P. To err (meiotically) is human: the genesis of human aneuploidy. Nature reviews. Genetics 2, 280–291, doi:10.1038/35066065 (2001).

3 Nagaoka, S. I., Hassold, T. J. & Hunt, P. A. Human aneuploidy: mechanisms and new insights into an age-old problem. Nature reviews. Genetics 13, 493–504, doi:10.1038/nrg3245 (2012).

4 Bongso, A., Chye, N. S., Ratnam, S., Sathananthan, H. & Wong, P. C. Chromosome anomalies in human oocytes failing to fertilize after insemination in vitro. Hum Reprod 3, 645–649, doi:10.1093/oxfordjournals.humrep.a136760 (1988).

5 Herbert, M., Kalleas, D., Cooney, D., Lamb, M. & Lister, L. Meiosis and maternal aging: insights from aneuploid oocytes and trisomy births. Cold Spring Harbor perspectives in biology 7, a017970, doi:10.1101/cshperspect.a017970 (2015).

6 Crawford, N. M. & Steiner, A. Z. Age-related infertility. Obstet Gynecol Clin North Am 42, 15–25, doi:10.1016/j.ogc.2014.09.005 (2015).

7 Lathi, R. B., Gray Hazard, F. K., Heerema-McKenney, A., Taylor, J. & Chueh, J. T. First trimester miscarriage evaluation. Semin Reprod Med 29, 463–469, doi:10.1055/s-0031-1293200 (2011).

8 Thomas, C., Cavazza, T. & Schuh, M. Aneuploidy in human eggs: contributions of the meiotic spindle. Biochem Soc Trans 49, 107–118, doi:10.1042/BST20200043 (2021).

9 Booth, D. G., Takagi, M., Sanchez-Pulido, L., Petfalski, E., Vargiu, G., Samejima, K., Imamoto, N., Ponting, C. P., Tollervey, D., Earnshaw, W. C. & Vagnarelli, P. Ki-67 is a PP1-interacting protein that organises the mitotic chromosome periphery. Elife 3, e01641, doi:10.7554/eLife.01641 (2014).

10 Strasburger, E. Ueber den Theilungsvorgang der Zellkerne und das Verhaltniss der Kerntheilung zur Zelltheilung. Arch Mikrosk Anat 21, 476–588 (1882).

11 Booth, D. G., Beckett, A. J., Molina, O., Samejima, I., Masumoto, H., Kouprina, N., Larionov, V., Prior, I. A. & Earnshaw, W. C. 3D-CLEM Reveals that a Major Portion of Mitotic Chromosomes Is Not Chromatin. Molecular cell 64, 790–802, doi:10.1016/j.molcel.2016.10.009 (2016).

12 Cuylen, S., Blaukopf, C., Politi, A. Z., Muller-Reichert, T., Neumann, B., Poser, I., Ellenberg, J., Hyman, A. A. & Gerlich, D. W. Ki-67 acts as a biological surfactant to disperse mitotic chromosomes. Nature 535, 308–312, doi:10.1038/nature18610 (2016).

13 Hernandez-Armendariz, A., Sorichetti, V., Hayashi, Y., Koskova, Z., Brunner, A., Ellenberg, J., Saric, A. & Cuylen-Haering, S. A liquid-like coat mediates chromosome clustering during mitotic exit. Mol Cell 84, 3254–3270 e3259, doi:10.1016/j.molcel.2024.07.022 (2024).

14 Sirri, V., Jourdan, N., Hernandez-Verdun, D. & Roussel, P. Sharing of mitotic pre-ribosomal particles between daughter cells. J Cell Sci 129, 1592–1604, doi:10.1242/jcs.180521 (2016).

15 Takagi, M., Natsume, T., Kanemaki, M. T. & Imamoto, N. Perichromosomal protein Ki67 supports mitotic chromosome architecture. Genes Cells 21, 1113–1124, doi:10.1111/gtc.12420 (2016).

16 Sobecki, M. et al. The cell proliferation antigen Ki-67 organises heterochromatin. eLife 5, e13722, doi:10.7554/eLife.13722 (2016).

17 Garwain, O., Sun, X., Iyer, D. R., Li, R., Zhu, L. J. & Kaufman, P. D. The chromatin-binding domain of Ki-67 together with p53 protects human chromosomes from mitotic damage. Proc Natl Acad Sci U S A 118, doi:10.1073/pnas.2021998118 (2021).

18 Booth, D. G. & Earnshaw, W. C. Ki-67 and the Chromosome Periphery Compartment in Mitosis. Trends Cell Biol 27, 906–916, doi:10.1016/j.tcb.2017.08.001 (2017).

19 Sun, X. & Kaufman, P. D. Ki-67: more than a proliferation marker. Chromosoma 127, 175–186, doi:10.1007/s00412-018-0659-8 (2018).

20 Remnant, L., Kochanova, N. Y., Reid, C., Cisneros-Soberanis, F. & Earnshaw, W. C. The intrinsically disorderly story of Ki-67. Open Biol 11, 210120, doi:10.1098/rsob.210120 (2021).

21 Gerdes, J., Schwab, U., Lemke, H. & Stein, H. Production of a mouse monoclonal antibody reactive with a human nuclear antigen associated with cell proliferation. Int J Cancer 31, 13–20, doi:10.1002/ijc.2910310104 (1983).

22 Traut, W., Endl, E., Scholzen, T., Gerdes, J. & Winking, H. The temporal and spatial distribution of the proliferation associated Ki-67 protein during female and male meiosis. Chromosoma 111, 156–164, doi:10.1007/s00412-002-0202-8 (2002).

23 Booth, D. G., Beckett, A. J., Prior, I. A. & Meijer, D. SuperCLEM: an accessible correlative light and electron microscopy approach for investigation of neurons and glia in vitro. Biol Open 8, doi:10.1242/bio.042085 (2019).

24 Cisneros-Soberanis, F., Simpson, E. L., Beckett, A. J., Pucekova, N., Corless, S., Kochanova, N. Y., Prior, I. A., Booth, D. G. & Earnshaw, W. C. Near millimolar concentration of nucleosomes in mitotic chromosomes from late prometaphase into anaphase. J Cell Biol 223, doi:10.1083/jcb.202403165 (2024).

25 Bonnet-Garnier, A., Feuerstein, P., Chebrout, M., Fleurot, R., Jan, H. U., Debey, P. & Beaujean, N. Genome organization and epigenetic marks in mouse germinal vesicle oocytes. Int J Dev Biol 56, 877–887, doi:10.1387/ijdb.120149ab (2012).

26 Verheijen, R., Kuijpers, H. J., Schlingemann, R. O., Boehmer, A. L., van Driel, R., Brakenhoff, G. J. & Ramaekers, F. C. Ki-67 detects a nuclear matrix-associated proliferation-related antigen. I. Intracellular localization during interphase. J Cell Sci 92 (Pt 1), 123–130, doi:10.1242/jcs.92.1.123 (1989).

27 McIntosh, J. R. & Landis, S. C. The distribution of spindle microtubules during mitosis in cultured human cells. J Cell Biol 49, 468–497, doi:10.1083/jcb.49.2.468 (1971).

28 Mosgoller, W., Leitch, A. R., Brown, J. K. & Heslop-Harrison, J. S. Chromosome arrangements in human fibroblasts at mitosis. Hum Genet 88, 27–33, doi:10.1007/BF00204924 (1991).

29 Takenouchi, O., Sakakibara, Y. & Kitajima, T. S. Live chromosome identifying and tracking reveals size-based spatial pathway of meiotic errors in oocytes. Science 385, eadn5529, doi:10.1126/science.adn5529 (2024).

30 Clift, D., So, C., McEwan, W. A., James, L. C. & Schuh, M. Acute and rapid degradation of endogenous proteins by Trim-Away. Nat Protoc 13, 2149–2175, doi:10.1038/s41596-018-0028-3 (2018).

31 Zuccotti, M., Ponce, R. H., Boiani, M., Guizzardi, S., Govoni, P., Scandroglio, R., Garagna, S. & Redi, C. A. The analysis of chromatin organisation allows selection of mouse antral oocytes competent for development to blastocyst. Zygote 10, 73–78, doi:10.1017/s0967199402002101 (2002).

32 Stenstrom, L., Mahdessian, D., Gnann, C., Cesnik, A. J., Ouyang, W., Leonetti, M. D., Uhlen, M., Cuylen-Haering, S., Thul, P. J. & Lundberg, E. Mapping the nucleolar proteome reveals a spatiotemporal organization related to intrinsic protein disorder. Mol Syst Biol 16, e9469, doi:10.15252/msb.20209469 (2020).

33 Ruebel, M. L., Schall, P. Z., Midic, U., Vincent, K. A., Goheen, B., VandeVoort, C. A. & Latham, K. E. Transcriptome analysis of rhesus monkey failed-to-mature oocytes: deficiencies in transcriptional regulation and cytoplasmic maturation of the oocyte mRNA population. Mol Hum Reprod 24, 478–494, doi:10.1093/molehr/gay032 (2018).

34 Akera, T., Chmatal, L., Trimm, E., Yang, K., Aonbangkhen, C., Chenoweth, D. M., Janke, C., Schultz, R. M. & Lampson, M. A. Spindle asymmetry drives non-Mendelian chromosome segregation. Science 358, 668–672, doi:10.1126/science.aan0092 (2017).

35 Sunchu, B. & Cabernard, C. Principles and mechanisms of asymmetric cell division. Development 147, doi:10.1242/dev.167650 (2020).

36 Duijf, P. H., Schultz, N. & Benezra, R. Cancer cells preferentially lose small chromosomes. Int J Cancer 132, 2316–2326, doi:10.1002/ijc.27924 (2013).

37 Zheng, J. N., Ma, T. X., Cao, J. Y., Sun, X. Q., Chen, J. C., Li, W., Wen, R. M., Sun, Y. F. & Pei, D. S. Knockdown of Ki-67 by small interfering RNA leads to inhibition of proliferation and induction of apoptosis in human renal carcinoma cells. Life Sci 78, 724–729, doi:10.1016/j.lfs.2005.05.064 (2006).

38 Ferrandiz, N., Downie, L., Starling, G. P. & Royle, S. J. Endomembranes promote chromosome missegregation by ensheathing misaligned chromosomes. J Cell Biol 221, doi:10.1083/jcb.202203021 (2022).

39 Yang, C., Zhang, J., Ding, M., Xu, K., Li, L., Mao, L. & Zheng, J. Ki67 targeted strategies for cancer therapy. Clin Transl Oncol 20, 570–575, doi:10.1007/s12094-017-1774-3 (2018).

